# The developing midbrain–hindbrain boundary contains molecularly distinct cell populations

**DOI:** 10.64898/2026.07.07.737085

**Authors:** Sylvia A. Nunez, Yong-Il Kim, Rebecca O’Rourke, Charles G. Sagerström

## Abstract

**Background:** During vertebrate embryogenesis, the isthmic region spans the midbrain–hindbrain boundary of the neural tube and includes an organizer (IsO) that is essential for proper formation of adjacent brain regions, yet the molecular and cellular composition of the isthmic region remains unresolved.

**Results:** We employed combined single-nucleus ATAC-seq and RNA-seq (scMultiome) in 13 and 16 hours-post-fertilization zebrafish embryos to molecularly resolve cell populations in the isthmic region and validated our findings in vivo by RNA fluorescence in situ hybridization. We identified two distinct isthmic cell populations – isthmic midbrain (IsMB) and isthmic hindbrain (IsHB) – that share expression of canonical isthmic genes, but that differ in their expression of midbrain vs hindbrain genes. We also uncovered a previously unrecognized heterogeneity within the IsHB, reflecting a canonical *fgf8*-expressing population anteriorly (IsO/r0a), and a novel *fgf8*-negative population posteriorly (r0p). We find that inhibition of Fgf signaling disrupts formation of the isthmic region, leading to loss of isthmic cell populations except a residual population characterized by a mixed neural identity.

**Conclusions:** Using transcriptional and epigenetic characterization, we expand on prior anatomical and genetic analyses of the isthmic region to refine our understanding of its cellular organization and demonstrate that it consists of several subdomains.

## INTRODUCTION

Embryonic development requires accurate organization, growth, and differentiation of cells into specific tissues and organs. Developmental signaling centers, known as organizers, are specialized groups of cells that secrete morphogens which influence the fate of surrounding cells and pattern the developing embryo. The organizer concept was first established through the classical Spemann-Mangold transplantation experiments, which demonstrated that a localized embryonic cell population could induce the formation of a secondary body axis and thereby coordinate surrounding tissue patterning (Spemann and Mangold, 1924; De Robertis, 2006; Arias and Steventon, 2018). Although the concept of organizers has been recognized for nearly a century, we still do not fully understand the cellular and molecular mechanisms by which they regulate development. Identifying how organizers are specified, how they interact with surrounding tissues, and how their signaling is integrated into developmental gene regulatory networks is essential for elucidating the principles that guide vertebrate development.

In the developing vertebrate central nervous system, regional identity is partly established through progressive subdivision of the neural tube into molecularly and morphologically distinct domains along the anterior–posterior axis. Early neuroanatomical studies described the compartmental organization of the neural tube into neuromeres, and later work established that the hindbrain is divided into rhombomeres, which behave as segmental units of gene expression, cell lineage restriction, and neuronal patterning (Orr, 1887; Vaage, 1969; Lumsden and Keynes, 1989; Kulesa and Fraser, 1998). At the interface between the midbrain and hindbrain lies a secondary neural organizer, the isthmic organizer (IsO), located at the isthmic constriction of the midbrain–hindbrain boundary (MHB). The IsO functions as a key signaling center that patterns the posterior midbrain and anterior hindbrain, including structures that contribute to the tectum and cerebellum (Alvarado-Mallart and Sotelo, 1984; Nakamura, 1990; Martínez et al., 1991; Wurst and Bally-Cuif, 2001; Liu and Joyner, 2001; Hidalgo-Sánchez et al., 2022). Recent reviews emphasize that the gene regulatory network of the isthmic region is deeply conserved and it remains a major model for understanding how signaling centers coordinate regional brain development (Hidalgo-Sánchez et al., 2022; Chandel and Hörnblad, 2025; Yuan et al., 2025).

The isthmic region spans the interface of two major transcriptional domains: anterior *Otx1/2* expression and posterior *Gbx1/2* expression. Mutual repression between Otx and Gbx factors positions the isthmic constriction and establishes a boundary at which organizer-associated genes are subsequently activated (Joyner, Liu, and Millet, 2000; Rhinn and Brand, 2001; Wurst and Bally-Cuif, 2001). Shortly thereafter, members of the Pax2/5/8 and Engrailed1/2 families become expressed in the isthmic region and contribute to organizer specification and maintenance. Pax2 is important for initiating the IsO program, including activation of *Fgf8*, the central signaling molecule of the organizer (Brand et al., 1996; Pfeffer et al., 1998; Ye et al., 2001; Ninkovic et al., 2005). The gene regulatory network of the isthmic region also includes Wnt1, Lmx1b, Irx1/2, and Fgf-responsive genes such as *Spry* and *ETS*-family targets, which together define the isthmic region and support organizer activity (Rhinn and Brand, 2001; Wurst and Bally-Cuif, 2001; Hidalgo-Sánchez et al., 2022; Chandel and Hörnblad, 2025).

Functional studies have demonstrated that Fgf signaling is essential for IsO activity. Ectopic Fgf8 can induce an ectopic organizer and produce mirror-image duplications of midbrain and cerebellar structures, demonstrating that Fgf8 is sufficient to confer organizer-like patterning activity (Crossley et al., 1996; Martínez et al., 1999). Conversely, loss of Fgf8 disrupts MHB development, affects *Wnt1* and *Gbx2* expression, and compromises survival of prospective midbrain and cerebellar tissues (Reifers et al., 1998; Chi et al., 2003). Fgf8 signaling also patterns the anterior hindbrain and helps establish the anterior limit of Hox gene expression, thereby contributing to rhombomere 1 (r1) identity (Irving and Mason, 2000). These studies place Fgf8 at the center of IsO function.

Despite extensive molecular characterization of the signaling pathways in the isthmic region, its cellular heterogeneity remains poorly resolved. The isthmic region is broadly defined by expression of *pax2/5/8* and *en1/2*, but this territory includes both posterior midbrain and anterior hindbrain domains, suggesting that distinct cell populations are likely to exist within the isthmic region. Further, the most anterior hindbrain segment, r1, is distinct from more posterior rhombomeres because it lacks Hox gene expression and gives rise to cerebellar structures (Irving and Mason, 2000; Wurst and Bally-Cuif, 2001). Accordingly, prior studies have proposed the existence of a distinct anterior hindbrain domain, sometimes referred to as r0 (Waskiewicz et al., 2002; Puelles, 2024). While this distinct r0 segment anterior to r1 has been predicted based on the expression domains of genes such as *fgf3r* and *ephA4a* (Walshe and Mason, 2000; Sleptsova-Friedrich, et al., 2001; Waskiewicz et al., 2002), it is not clear if this distinction is supported by additional molecular characteristics. Nor is it clear how the *fgf8*-expressing IsO relates to the rest of the isthmic cells in molecular terms (Harada et al., 2016; Hidalgo-Sánchez et al., 2022).

Determining whether the isthmic region is homogeneous or composed of multiple cell populations is essential for refining models of brain development, particularly in terms of the IsO.

Recent advances in single-cell technologies have made it possible to characterize the molecular state of individual cells in the isthmic region. Single-cell multiome (scMultiome) approaches simultaneously capture gene expression and chromatin accessibility, allowing transcriptional identity to be linked with candidate regulatory programs in individual cells. In zebrafish, our recent scMultiome work has shown that hindbrain rhombomeres become molecularly distinct between the end of gastrulation and early segmentation stages, providing a framework for resolving cell states during early brain regionalization (Kim et al., 2023; Warns et al., 2024). When combined with spatial validation methods such as hybridization chain reaction RNA fluorescence in situ hybridization (HCR RNA-FISH), these approaches enable molecularly defined cell populations to be mapped back onto the developing embryo.

Here, we use scMultiome analysis together with HCR RNA-FISH to examine cellular heterogeneity within the developing zebrafish isthmus at 13 and 16 hours-post-fertilization (hpf). Using *pax* and *engrailed* gene expression to identify isthmic cells, we detect two major cell populations: an isthmic midbrain population (IsMB), that is defined by also expressing midbrain-associated genes such as *otx1/2* and *wnt1*, and an isthmic hindbrain population (IsHB), that co-expresses hindbrain-associated genes including *gbx2* and *fgf8*. We show that the IsHB is molecularly distinct from r1, and we predict that it corresponds to the previously hypothesized r0. We further find that IsHB/r0 consists of two molecularly distinct subdomains: an *fgf8*-expressing population that we refer to as anterior r0 (r0a; and that likely corresponds to the canonical IsO) and a previously undescribed *fgf8*-negative population, which we refer to as posterior r0 (r0p). Finally, by inhibiting Fgf signaling, we demonstrate that formation of these isthmic subpopulations depends on Fgf activity. Our findings refine the traditional model of the vertebrate isthmus region by revealing multiple molecularly distinct cell populations.

## RESULTS

### The zebrafish isthmic region can be divided into isthmic midbrain and isthmic hindbrain

To examine heterogeneity of the isthmus region during early embryonic development, we analyzed our previously published scMultiome data (Kim et al., 2023) collected at 16hpf – a time corresponding to mid-segmentation stages, when the isthmus is first becoming morphologically observable in zebrafish (Gutzman et al., 2008). Using canonical gene markers for the isthmic region, we identified two isthmic populations in the 16hpf UMAP (Fig. 1A – black arrows). Both populations express *en2a, en2b, pax2a, pax2b* and *pax5* (Fig. 1B) – genes that are known to define the isthmic region. Further analysis of differentially expressed genes (DEGs) revealed that one of these isthmic populations co-expresses genes involved in midbrain formation (e.g., *otx1*, *otx2a* and *otx2b;* Fig. 1B). Upon closer examination, we also find that this population expresses *wnt1*, which defines the posterior midbrain adjacent to the isthmus (Fig. 1C). We conclude that this isthmic population has midbrain characteristics and we refer to it as isthmic midbrain (IsMB). A second isthmic population is located between the IsMB and rhombomere 1 (r1) in the UMAP (Fig. 1A) and shows significant enrichment for the expression of *gbx2* (Fig. 1D), which is known to regulate hindbrain formation. This population also expresses *fgf8a* and *fgf8b* (Fig. 1B, 1D) that are known markers of the anterior hindbrain adjacent to the isthmus. Since this population displays hindbrain characteristics, we refer to it as isthmic hindbrain (IsHB). We note that the IsHB population forms a distinct cluster in the UMAP, while the IsMB population maps closely to midbrain cells. To further assess if the IsMB and IsHB populations are distinct from each other, and from the adjacent midbrain (MB) and hindbrain (r1) populations, we generated heatmaps showing the expression of the top DEGs for the IsMB and the IsHB (Fig. 1C, D, respectively) across all four clusters. This analysis reveals that the IsMB and IsHB populations are transcriptionally distinct from midbrain and r1 cells. These identities are further supported by our identification of PAX transcription factor binding motifs as being enriched in accessible chromatin regions in both the IsMB (Fig. 1E, 1G) and the IsHB (Fig. 1F, 1H) populations, while motifs for midbrain-restricted genes (e.g., OTX and DMBX) are enriched in the IsMB, but not the IsHB. Taken together, these transcriptomic and chromatin accessibility profiles indicate that the 16hpf zebrafish isthmus contains two molecularly distinct populations corresponding to IsMB and IsHB territories.

**Figure 1.**
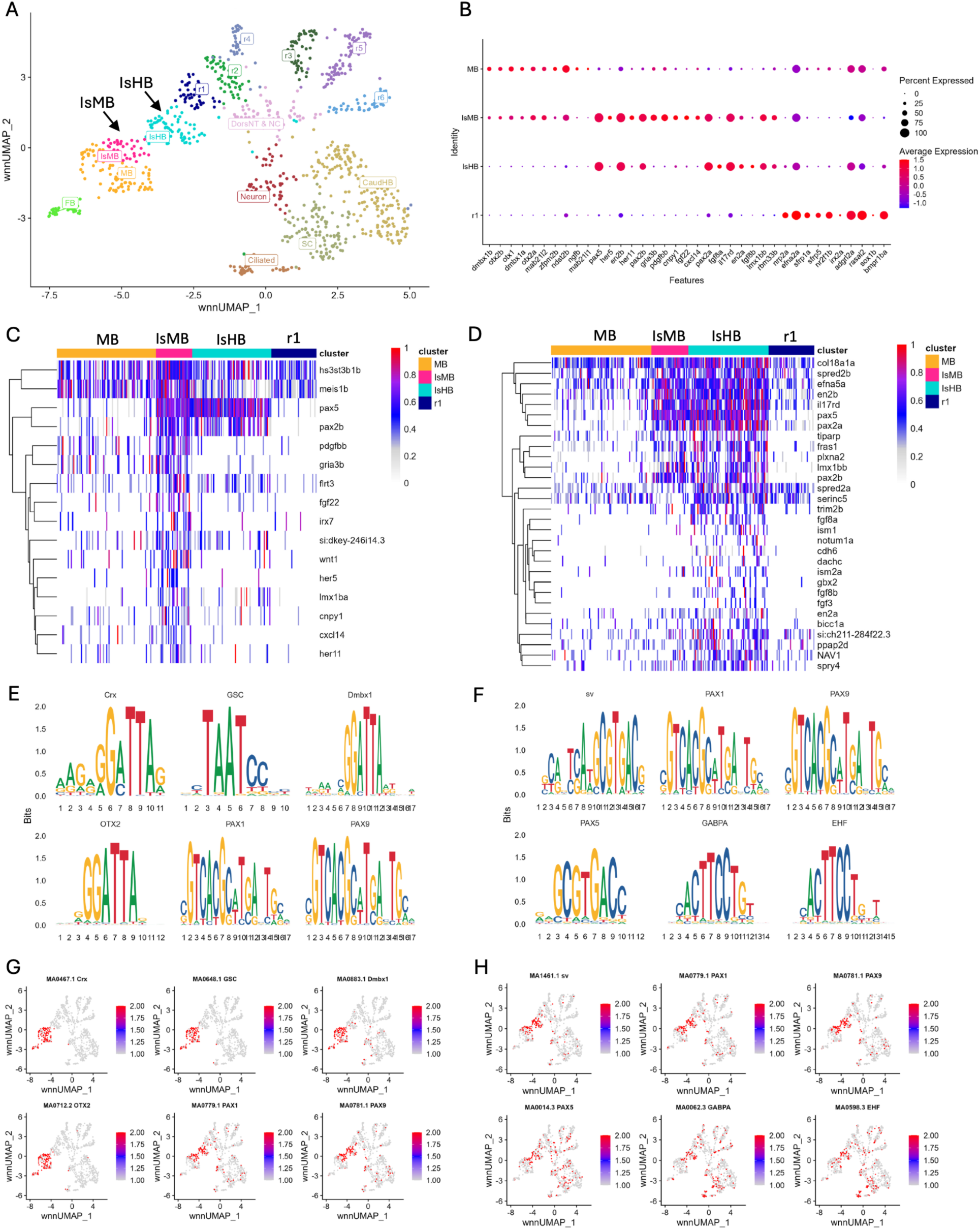
Two subpopulations can be molecularly resolved in the 16hpf zebrafish isthmic region. (A) UMAP of 16hpf neural clusters. (B) Dot plot showing expression of top ten enriched genes in the MB, IsMB, IsHB, and r1 clusters. (C, D) Heatmaps showing expression of top DEGs derived from IsMB (C) or IsHB (D). (E-H). Six selected enriched binding motifs in the IsMB (E, G) and IsHB (F, H) subpopulations shown as motif logo (E, F) or UMAP (G, H). For reference, the IsMB cluster defined here corresponds to one cluster (MHB.2), and the IsHB corresponds to two clusters (MHB.1 and MHB.3), that we provisionally identified, but did not characterize, at 16hpf previously (Kim et al., 2023). Abbreviations: CaudHB = caudal hindbrain, DEG = differentially expressed gene, FB = forebrain, HB = hindbrain, Is = isthmus, IsHB = isthmus hindbrain, IsMB = isthmus midbrain, MB = midbrain, MHB = midbrain–hindbrain boundary, r = rhombomere, SC = spinal cord, UMAP = Uniform Manifold Approximation and Projection.

To determine if the IsMB and IsHB populations defined by our scMultiome analysis are present in the intact embryo in vivo, we next used HCR to assess gene expression in 16hpf zebrafish embryos. Two gene markers for the isthmic region – *en2b* (Fig. 2A, D, J, M, W) and *pax2a* (Fig. 2G, Z) – robustly delineate the isthmic region and positions it between two *nr2f2* expression domains (Ghosh, et al., 2018; Love and Prince, 2012) that mark the midbrain anteriorly and r1 posteriorly (Fig. 2A-C, W, X; bracket in C marks the isthmic region). Co-detection of *en2b* together with *otx1* – that marks the anterior embryo, including the midbrain – revealed partial overlap such that approximately the anterior half of the *en2b* expression domain overlaps with *otx1* expression (Fig. 2D-F, W, Y). This pattern coincides with the overlap between these two markers observed in the IsMB population in our scMultiome analysis, and we conclude that this co-expression domain corresponds to the IsMB in vivo (bracket in Fig. 2F). In further support of the IsMB being present in vivo, we also find that expression of the *pax2a* isthmus marker overlaps with expression of *wnt1* – that marks the posterior midbrain (Fig. 2G-I, Z, AA; bracket in I marks *pax2a/wnt1* overlap). In an analogous manner, *en2b* partially overlaps with the hindbrain markers *gbx2* (Fig. 2J-L, W, BB) and *fgf8a* (Fig. 2M-O, W, FF) posteriorly, identifying the IsHB in vivo (brackets in Fig. 2L, O). To further assess the molecular distinction between the IsMB and the IsHB in vivo, we selected one marker for each (*her5* for IsMB and *ism1* for IsHB) from our DEG analysis in Fig. 1B-D. Co-detection of these markers with the *en2a* isthmic marker shows that *her5* (Fig. 2S-V, CC, EE) and *ism1* (Fig. 2S-V, DD, EE) both overlap with *en2a* expression, but not with each other. Together these data are consistent with distinct IsMB and IsHB populations existing as spatially and molecularly distinct domains within the developing zebrafish isthmus in vivo.

**Figure 2.**
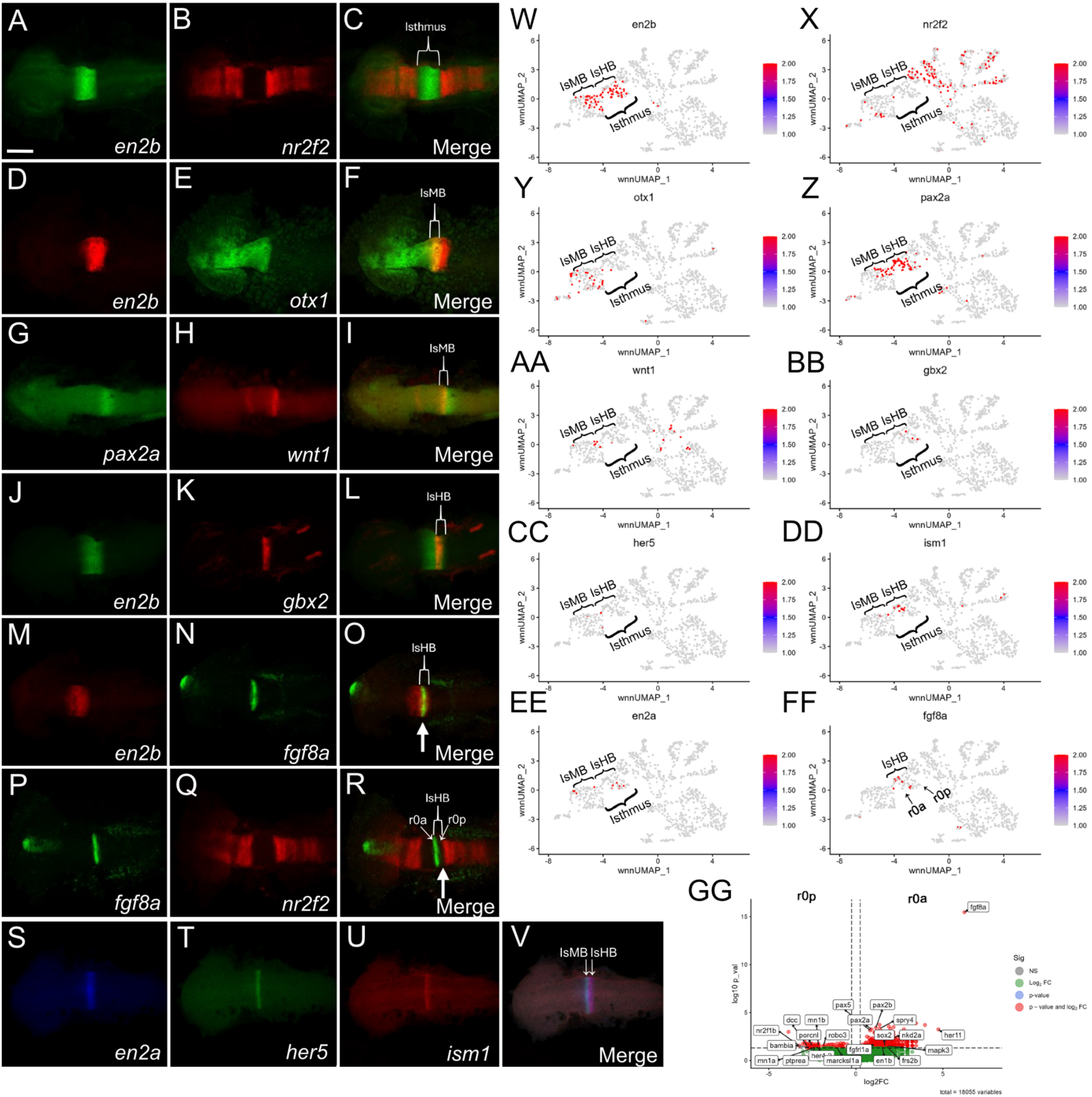
The IsMB and IsHB subpopulations are present as distinct domains in 16hpf zebrafish. (A-V) HCR was used to detect expression of the gene indicated at the bottom of each panel (C, F, I, L, O, R and V represent merged images of the panels to their left). Panels show flat-mounted 16hpf zebrafish embryos in dorsal view with anterior to the left. Scale bar = 100 um. White arrow in R indicates gap between *fgf8a* and *nr2f2* expression. (W-GG) Feature plots showing cells in the UMAP that express the gene indicated at the top of each panel. Cell populations discussed in the text are indicated in each panel. (GG) Volcano plot of DEGs identified by comparing *fgf8*-expressing (r0a) vs. *fgf8-*negative cells (r0p) in the IsHB.

### The isthmic hindbrain consists of *fgf8a*-expressing and *fgf8a*-negative domains

Because *fgf8a* expression is known to mark the isthmic organizer (IsO), and we detect *fgf8* expression in the IsHB, it is possible that the IsHB corresponds to the IsO. To test this possibility, we examined *fgf8a* expression in greater detail. Our HCR analyses revealed that *fgf8* expression (arrow in Fig. 2O) is restricted to a subset of IsHB cells and that *fgf8a* expression does not extend all the way posteriorly to the *nr2f2* expression domain that marks r1 (Fig. 2P-R; arrow in panel R indicates gap between *fgf8a* expression and r1), suggesting that the IsHB can be subdivided further. Examining *fgf8a* expression in the UMAP also revealed distinct populations of *fgf8a*-expressing cells, which we refer to as anterior r0 or r0a, and *fgf8a*-negative cells, which we refer to as posterior r0 or r0p, in the IsHB (Fig. 2FF). Subclustering of the IsHB based on *fgf8a* expression, followed by displaying DEGs as a volcano plot, showed that *fgf8a* is the primary significant marker distinguishing these two subpopulations (Fig. 2GG). In addition to *fgf8a*, the *fgf8a*-expressing IsHB population showed enrichment of genes associated with FGF/MAPK signaling and feedback, including *spry4*, *fgfrl1a*, *frs2b*, and *mapk3*, as well as MHB/isthmic and neural progenitor-associated genes such as *pax2a*, *pax2b*, *pax5*, *en1b*, *her11*, *sox2*, and *nkd2a*. These nominally enriched genes support the interpretation that *fgf8a*-expressing IsHB cells (r0a) represent an FGF-active, organizer-like neural progenitor state at 16hpf. In contrast, the *fgf8a*-negative IsHB cells do not show a strong canonical MHB/IsO organizer signature. Instead, their enrichment of genes suggests a more neural-patterned or differentiating state, with enrichment of axon-guidance/morphogenesis genes such as *dcc*, *robo3*, and *marcksl1a*, regional/transcriptional regulators such as *nr2f1b*, *her4.2*, *mn1a*, and *mn1b*, and signaling modulators such as *porcnl*, *bambia*, and *ptprea*. This may indicate that the *fgf8a*-negative IsHB population (r0p) is less organizer-like and more reflective of adjacent hindbrain/neural identity or a transitional neural state. When we apply a more stringent adjusted-p-value correction, only *fgf8a* expression in r0a scores as being enriched, demonstrating that r0a and r0b are highly related molecularly.

Collectively, these data indicate that the IsHB is not homogeneous, but that it itself consists of at least two subpopulations: an anterior *fgf8a*-expressing population that we refer to as r0a, and that likely corresponds to the IsO, and a posterior *fgf8a*-negative one that we refer to as r0p (Fig. 2R, FF).

### The isthmic region is subdivided by early segmentation stages

To establish if the IsMB and IsHB populations are present earlier during formation of the isthmic region, we next analyzed our scMultiome data obtained from 13hpf zebrafish. At this stage, the isthmus is not yet apparent morphologically, but subdomains have been proposed within the isthmic region based on gene expression patterns (Watson, et al. 2017; Puelles, 2024). We find the organization of the 13hpf UMAP to resemble that at 16hpf, as it includes two isthmic clusters (arrows in Fig. 3A). As at 16hpf, one of these clusters expresses midbrain markers, such as *otx1* and *wnt1,* alongside the isthmus markers *en2b* and multiple *pax* genes (*pax2a, pax5, pax7a;* Fig. 3B, C), identifying it as the IsMB. The other isthmic cluster expresses hindbrain markers, such as *fgf8b*, together with the isthmus markers *en2b* and *pax5* (Fig. 3B, D), defining it as the IsHB. Additionally, accessible chromatin regions in the IsMB and IsHB populations are enriched in binding motifs for isthmus-related transcription factors, including PAX proteins, and OTX motifs are enriched in the IsMB (Fig. 3E, G) but not the IsHB (Fig. 3F, H) populations. We also note that both the IsMB and IsHB populations are well resolved as distinct clusters at 13hpf, as opposed at 16hpf when the IsMB appeared more similar to the midbrain. We conclude that the IsMB and the IsHB are molecularly distinct by 13hpf in zebrafish.

**Figure 3.**
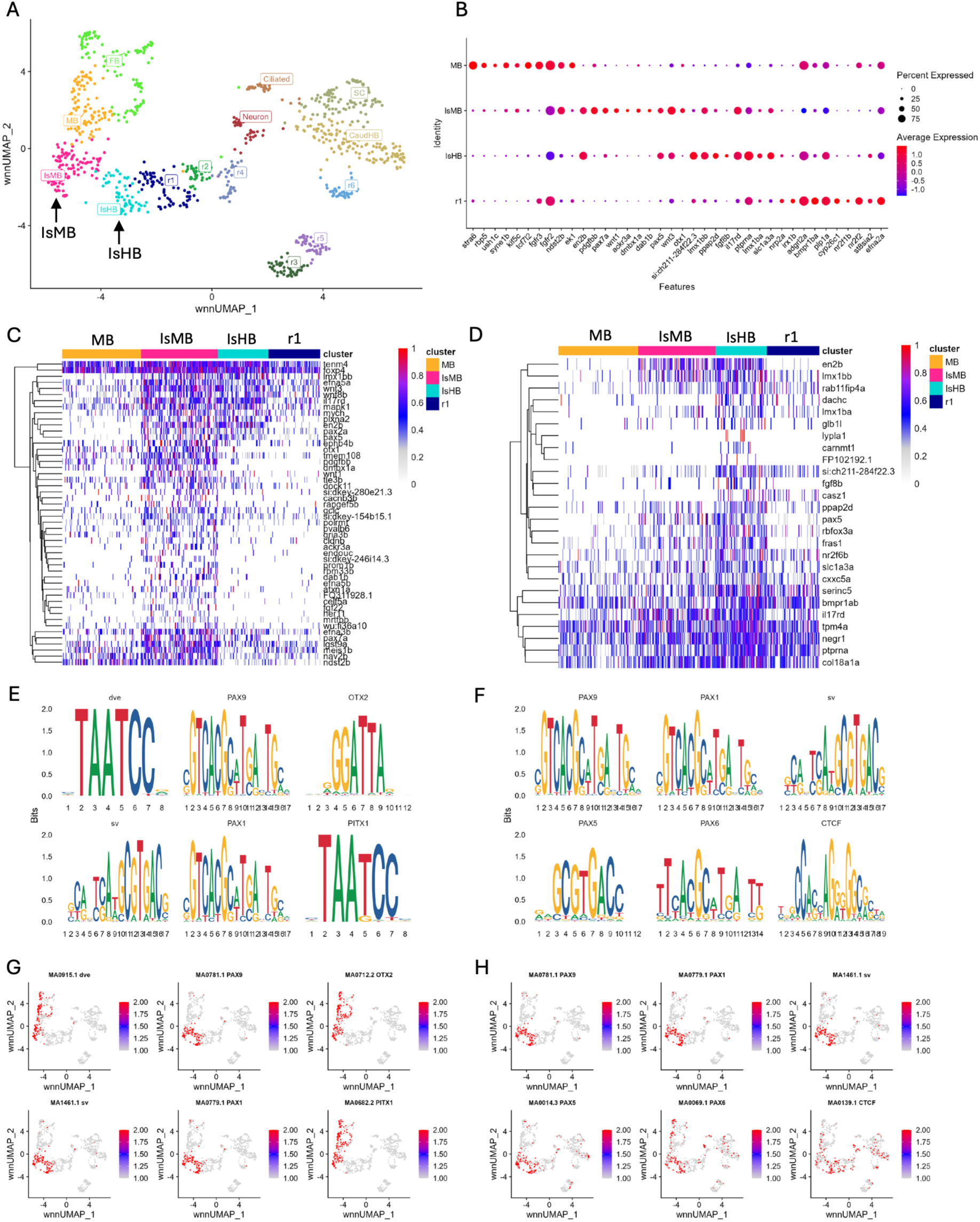
The IsMB and IsHB populations are present in the 13hpf UMAP. (A) UMAP of 13hpf neural clusters. (B) Dot plot showing expression of top ten enriched genes in MB, IsMB, IsHB, and r1 clusters. (C, D) Heatmaps showing expression of top DEGs derived from IsMB (C) or IsHB (D). (E-H) Six selected enriched binding motifs in the IsMB (E, G) and IsHB (F, H) subpopulations shown as motif logo (E, F) or UMAP (G, H). For reference, the IsMB cluster defined here corresponds to three clusters (MHB.2, MHB.3 and MHB.4), and the IsHB corresponds to two clusters (MHB.1 and MHB.5), that we provisionally identified, but did not characterize, at 13hpf previously (Kim et al., 2023). See legend to Figure 1 for abbreviations.

Using HCR analysis, we next tested if the IsMB and IsHB domains are also present in vivo already at 13hpf. This analysis revealed the presence of distinct IsMB and IsHB clusters, similar to the situation at 16hpf. The *en2b* isthmus marker spans the entirety of the isthmic region, as demonstrated by lack of overlap with *nr2f2* (Fig. 4A-C, S, T; bracket in C). *en2b* overlaps with the *otx1* midbrain marker (Fig. 4D-F, S, U), and the *pax2a* isthmus marker overlaps with *wnt1* (Fig. 4G-I, V, W), to define the IsMB (brackets in Fig. 4F, I), confirming that the IsMB is already present at 13hpf. In turn, the IsHB is identified by the co-localization of *en2b* and *gbx2* expression (Fig. 4J-L, S, X), in the posterior of the isthmic region (bracket in Fig. 4L marks the IsHB). As at 16hpf, *fgf8a* expression does not extend to the posterior-most edge of *en2b* expression (Fig. 4M-O, S, Y; arrow in panel O indicates *fgf8* expression) or to the anterior-most edge of *nr2f2* expression in r1 (Fig. 4P-R, T, Y; arrow in panel R indicates *fgf8*-negative cells). This analysis identified one population of *fgf8*-expressing cells (r0a, likely corresponding to the IsO) and one population of *fgf8*-negative cells (r0p) in the IsHB of 13hpf embryos (Fig. 4Z). We note that while these domains are readily observed in the embryo, they are less well resolved in the 13hpf UMAP, as *gbx2* and *fgf8* transcripts are barely detectable in the UMAP. Jointly, these results indicate that the isthmus subpopulations we identify, the IsMB and the IsHB – including the IsO/r0a and the novel r0p – are detectable at 13hpf both computationally and as spatially distinct domains in vivo, demonstrating the early establishment of isthmic substructure.

**Figure 4.**
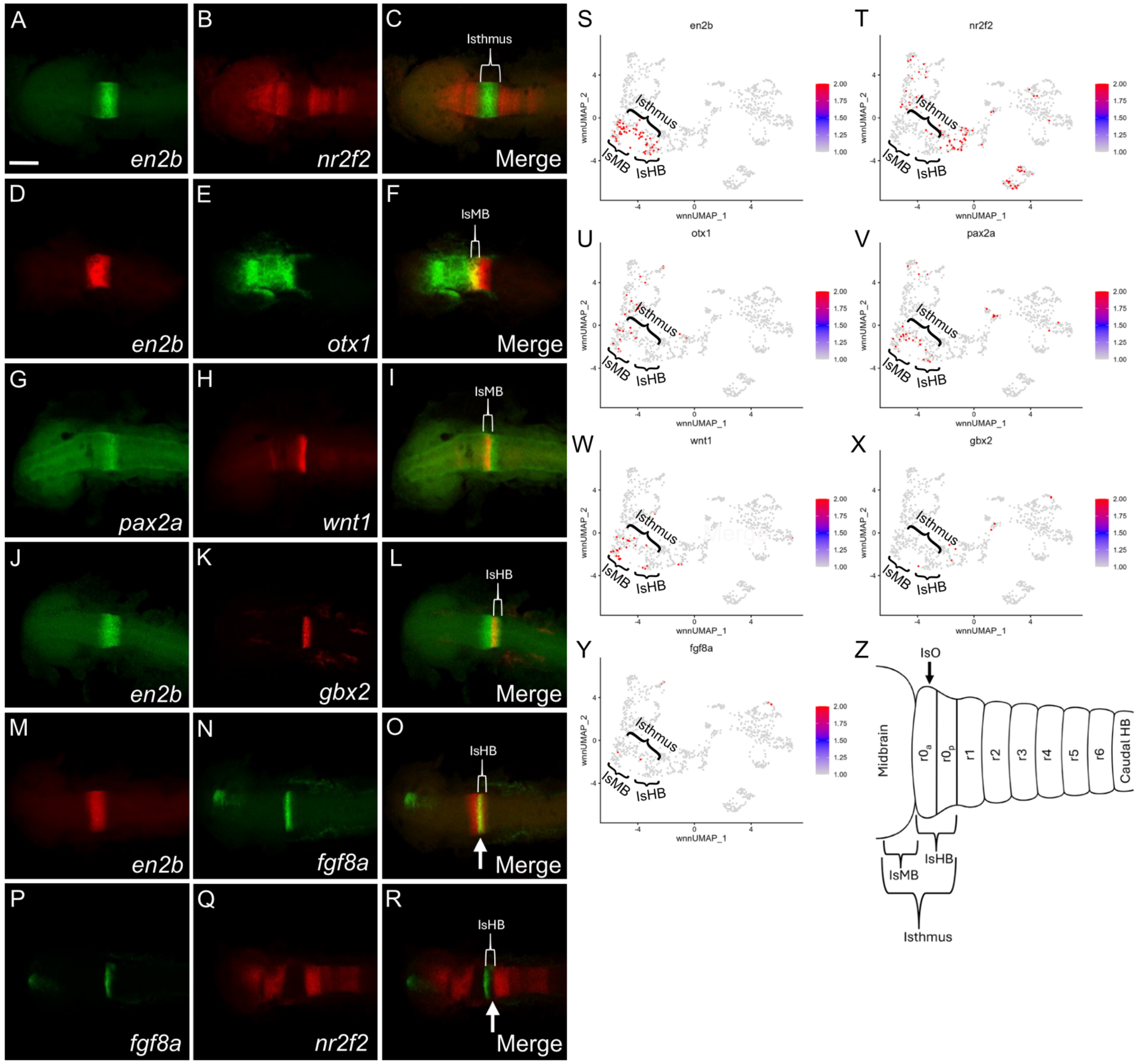
The IsMB and IsHB subpopulations are present in 13hpf zebrafish embryos. (A-R) HCR was used to detect expression of the gene indicated at the bottom of each panel (C, F, I, L, O and R represent merged images of the two panels to their left). Panels show flat-mounted 13hpf zebrafish embryos in dorsal view with anterior to the left. Scale bar = 100 um. White arrows indicate overlap between *en2b* and *fgf8a* expression (O) and gap between *fgf8a* and *nr2f2* expression (R). (S-Y) Feature plots showing cells in the UMAP that express the gene indicated at the top of each panel. Cell populations discussed in the text are indicated in each panel. (Z) Schematic diagram of subpopulations observed within the isthmic region.

### Formation of the IsMB and IsHB domains requires Fgf signaling

Fgf signaling is essential for formation and function of the isthmic region (Chi et al., 2003; Chandel and Hörnblad, 2025). Our identification of distinct subdomains within this region raises the question if formation of the IsMB and IsHB domains is dependent on Fgf signaling. To test this, we treated embryos with the Fgf receptor inhibitor SU5402, a small-molecule inhibitor of FGFR tyrosine kinase activity (originally characterized as an FGFR1 inhibitor (Mohammadi et al., 1997) and widely used to block FGF signaling in zebrafish embryos) and collected them for scMultiome analysis at 13hpf. scMultiome data from SU5402-treated embryos were reference-mapped to a 13hpf control dataset that we published previously (Kim et al., 2023) and then projected as a separate UMAP (Fig. 5A, B). For greater resolution in examining the effects of Fgf inhibition, we computationally subdivided the isthmic region further, which resulted in the IsMB cluster dividing into three clusters (MHB.2, MHB.3 and MHB.4) and the IsHB into two clusters (MHB.1 and MHB.5). An examination of the resulting UMAPs revealed a pronounced reduction in isthmic subpopulations in SU5402-treated embryos, with only a single distinct isthmus-related cluster remaining (arrow in Fig. 5B). Accordingly, comparing the fraction of cells that contribute to each neural cluster revealed that all clusters between the midbrain and the spinal cord are reduced in SU5402-treated embryos relative to control embryos. In the hindbrain, r2 and r5 appear lost, while r1, r3 and r4 are reduced, and at the isthmus only one main cluster, MHB.2, remains (Table 1). Our findings are consistent with previous reports demonstrating a requirement for Fgf signaling in the formation of the isthmic region, as well as of the hindbrain and the spinal cord, but also reveal an apparently Fgf-independent isthmic population.

**Figure 5.**
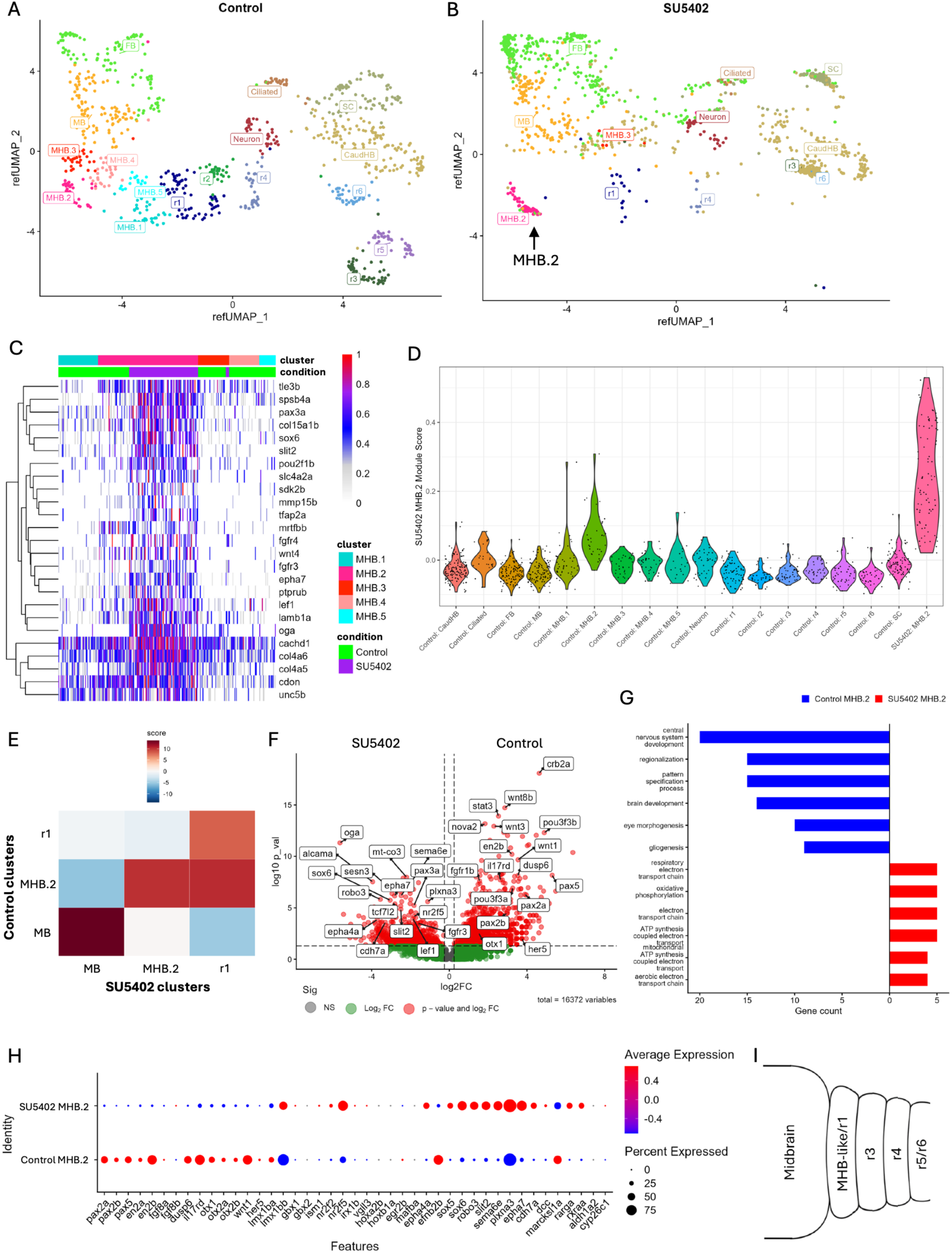
One predominant isthmic cell cluster remains after disruption of the FGF signaling pathway. (A, B) scMultiome data from 13hpf SU5402-treated embryos (B) were reference-mapped to 13hpf control embryos (A) and displayed as separate UMAPs. (C) Heatmap showing expression of top DEGs derived from the MHB.2 cluster in both control and SU5402-treated embryos. (D) A gene signature module generated from the remaining MHB.2 cluster in SU5402-treated embryos was used to derive a ModuleScore for each cluster in control embryos. (E) Heatmap showing Similarity Scores between selected cell clusters from control and SU5402-treated embryos. (F) Volcano plot displaying DEGs between MHB.2 clusters in control vs. SU5402-treated embryos. (G) Relative enrichment for GO terms in DEGs between MHB.2 clusters in control vs. SU5402-treated embryos. (H) Dot plot of selected genes between MHB.2 clusters in control vs. SU5402-treated embryos. (I) Schematic diagram of subpopulations that persist at the isthmic region following disruption of Fgf signaling. For reference, the MHB.1 – MHB.5 clusters defined here correspond to the MHB clusters that we provisionally identified (Kim et al., 2023), but did not characterize, at 13hpf previously.

**Table 1.** Cell counts for UMAP clusters in control and SU5402-treated 13hpf neural clusters.

| Cluster | Control (# cells) | Control (% cells) | SU5402 (# cells) | SU5402 (% cells) | Ratio (SU5402/Control) |
| --- | --- | --- | --- | --- | --- |
| FB | 111 | 11.7 | 354 | 28.0 | 3.19 |
| MB | 107 | 11.3 | 147 | 11.6 | 1.37 |
| MHB.1 | 46 | 4.8 | 0 | 0.0 | 0 |
| MHB.2 | 36 | 3.8 | 80 | 6.3 | 2.22 |
| MHB.3 | 32 | 3.4 | 4 | 0.3 | 0.12 |
| MHB.4 | 35 | 3.7 | 0 | 0.0 | 0 |
| MHB.5 | 19 | 2.0 | 0 | 0.0 | 0 |
| r1 | 71 | 7.5 | 19 | 1.5 | 0.27 |
| r2 | 35 | 3.7 | 0 | 0.0 | 0 |
| r3 | 47 | 4.9 | 6 | 0.5 | 0.13 |
| r4 | 38 | 4.0 | 8 | 0.6 | 0.21 |
| r5 | 43 | 4.5 | 0 | 0.0 | 0 |
| r6 | 36 | 3.8 | 1 | 0.1 | 0.03 |
| CaudHB | 144 | 15.2 | 404 | 31.9 | 2.81 |
| SC | 89 | 9.4 | 203 | 16.0 | 2.28 |
| Ciliated | 25 | 2.6 | 8 | 0.6 | 0.32 |
| Neuron | 36 | 3.8 | 31 | 2.5 | 0.86 |

To characterize the residual isthmic cell population in SU5402 treated embryos, we projected the top genes from this cluster in a heatmap across all isthmic clusters (Fig. 5C). In agreement with the UMAPs in Fig. 5A, B, the remaining MHB.2 cluster showed elevated expression of many genes expressed in the control MHB.2 cluster, such as *pax3a*, *wnt4*, and *epha7*. Despite this similarity, the MHB.2 cluster in SU5402-treated embryos, does not appear identical to the control MHB.2 cluster. To examine this in more detail, we generated a gene signature module score from the SU5402-treated MHB.2 population and assessed its similarity to each individual control neural cluster (Fig. 5D). This analysis found the control MHB.2 cluster as having the highest module score, indicating that the Fgf-independent MHB.2 population is most similar to the control MHB.2 state. Finally, a cluster similarity analysis supported this relationship (Fig. 5E), again identifying control MHB.2 as the closest match to the MHB.2 cluster from SU5402-treated embryos. We note that all three of these analyses (Fig. 5C-E) also demonstrate that these two populations are not identical. Of further note, the r1 cluster from SU5402-treated embryos showed stronger transcriptional similarity to control MHB.2 than to control r1, suggesting that the r1 population present after Fgf inhibition may be transcriptionally shifted toward an MHB.2-like state. These findings support the idea that Fgf signaling is required not only for maintaining the full complement of isthmic subdomains, but also for preserving normal r1 identity adjacent to the isthmus.

To characterize their differences, we identified DEGs between MHB.2 cluster cells from control and SU5402-treated embryos and projected them in a volcano plot. This analysis revealed that several canonical MHB-associated genes, including *wnt1*, *en2b*, *pax5*, and *pax2a*, were significantly reduced in SU5402-treated MHB.2 cells relative to control MHB.2 cells (Fig. 5F). In contrast, SU5402 MHB.2 cells showed significant enrichment of genes associated with neural adhesion and axon guidance, including *alcama*, *robo3*, *sema6e*, *epha7*, and *plxna3*, as well as genes associated with altered transcriptional or cellular state, such as *sox6*, *sesn3*, and *oga*. Additional significantly enriched genes included mitochondrial, cytoskeletal, and extracellular matrix-associated genes, suggesting broader changes in cellular physiology rather than maintenance of a canonical MHB program. This is consistent with our Gene Ontology (GO) term enrichment analysis of adjusted p-value significant MHB.2 DEGs, which revealed divergent top biological processes between Control and SU5402 conditions (Fig. 5G). Control MHB.2 cells were enriched for neurodevelopmental and patterning programs, including CNS development, regionalization, pattern specification, and brain development. Alternatively, SU5402 MHB.2 cells were enriched for mitochondrial respiration and oxidative phosphorylation pathways, suggesting that Fgf inhibition disrupts normal MHB developmental identity. These data indicate that while an MHB.2-related population persists after SU5402 treatment, it lacks key components of the normal isthmic gene expression program and instead adopts a transcriptionally altered MHB-like state following Fgf inhibition.

A dot plot of selected regional and differentially expressed genes further supported the altered identity of the residual SU5402 MHB.2 population (Fig. 5H). Control MHB.2 cells showed stronger expression of canonical MHB/isthmic organizer genes, including *pax2a*, *pax2b*, *pax5*, *en2a*, *en2b*, *wnt1*, *fgf8b*, *dusp6*, *il17rd*, and *her5*. By contrast, SU5402 MHB.2 cells showed increased expression of genes associated with neural differentiation, cell adhesion, axon guidance, and altered regional identity, including *sox6*, *robo3*, *slit2*, *sema6e*, *plxna3*, *epha7*, *cdh7a*, *rarga*, and *aldh1a2*. These data further indicate that the remaining SU5402 MHB.2 population is not equivalent to control MHB.2 but instead represents an MHB-like population with reduced isthmic organizer identity and increased expression of alternative neural-patterning genes.

Together, these analyses suggest that domains between the midbrain and the spinal cord are affected by disruption of Fgf-signaling, with many of these domains appearing to be lost. Only one Fgf-independent cluster is observed in SU5402-treated embryos and, while it is most similar to control MHB.2 cells, it does not fully retain the molecular identity of the normal MHB.2 population, leading us to designate it as MHB-like (Fig. 5I).

### Disruption of Fgf-signaling interferes with formation of the IsMB and the IsHB in vivo

To determine how the effects of disrupting Fgf-signaling that we observed by scMultiome analysis are reflected in vivo, we performed HCR on SU5402-treated embryos at 13hpf. We first examined the expression of canonical isthmic markers and find that expression of both *pax2a* and *en2b* is lost in SU5402-treated embryos (Fig. 6A, E, I, M). Using probes for *otx1*, *wnt1, gbx2*, and *fgf8a* – that mark the midbrain, the IsMB, the IsHB, and the IsO, respectively (Fig. 6Q-FF) – we find that SU5402-treated embryos display an expansion of the *otx1* domain (arrows in Fig. 6U and CC), and a loss of *gbx2*, *fgf8a*, and *wnt1* expression, relative to control embryos (Fig. R, S, V, W, AA, EE). These findings are consistent with the loss of isthmic clusters observed in the UMAP of SU5402-treated embryos.

**Figure 6.**
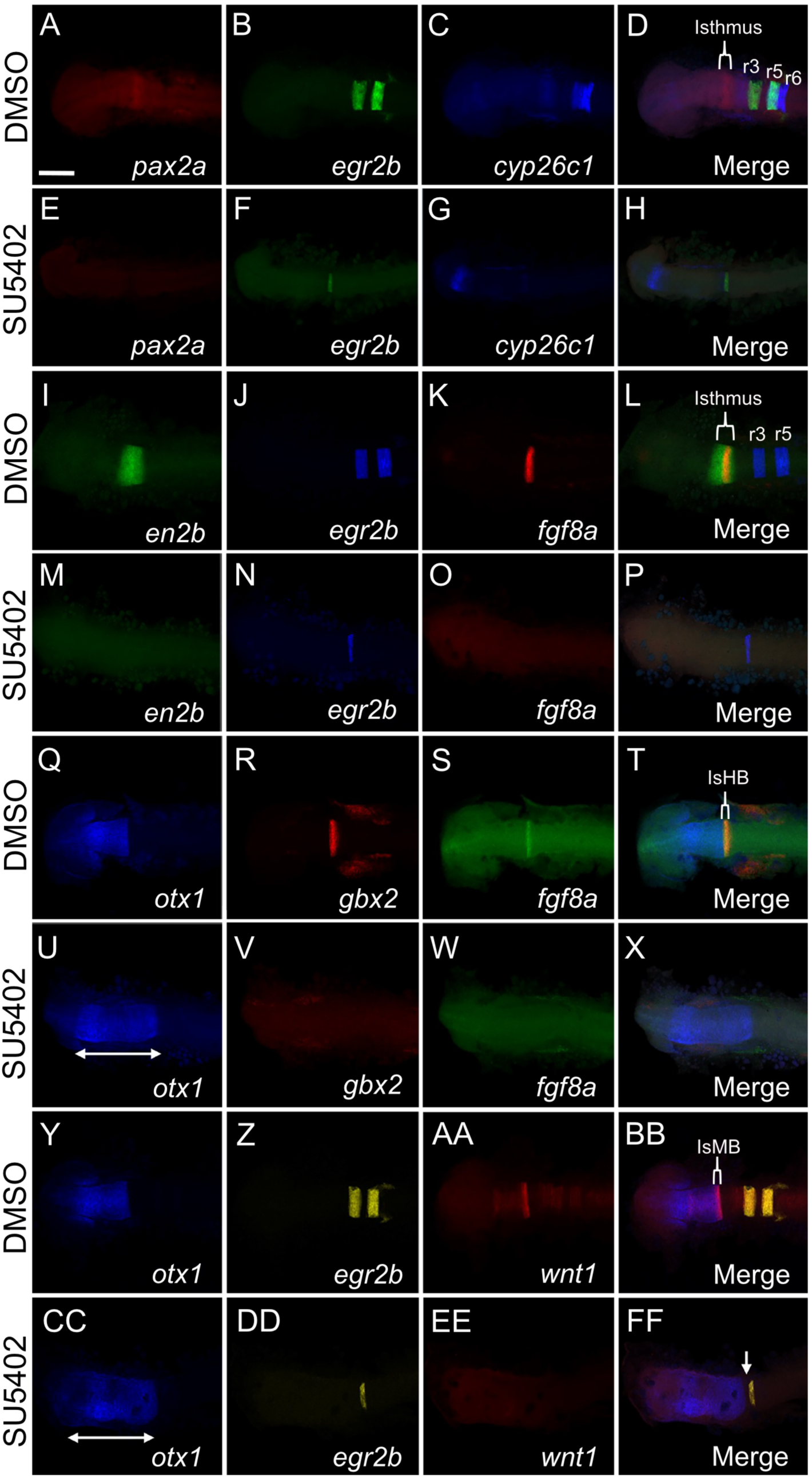
Isthmus and hindbrain development are severely impacted by disrupted Fgf signaling. HCR was used to detect expression of the gene indicated at the bottom of each panel (D, H, L, P, T, X, BB and FF represent merged images of the three panels to their left). Panels show flat-mounted 13hpf DMSO-treated (control) or SU5402-treated, as indicated at left, zebrafish embryos in dorsal view with anterior to the left. Scale bar = 100 um. White arrow in FF indicates gap between *otx1* and *egr2b* expression.

Because SU5402 treatment also disrupted hindbrain-associated clusters in the scMultiome analysis, we also examined hindbrain patterning in vivo. We find that expression of *egr2b* (that marks rhombomeres r3 and r5), *irx1b* (that marks r1), *vgll3* (that marks r2) and *mafba* (that marks r5 and r6) is disrupted in SU5402-treated embryos (Fig. 7). Expression of *vgll3* is lost (Fig. 7K, O), indicating that r2 is lost upon SU5402-treatment, in agreement with our scMultiome analysis (Fig. 5B, Table 1). *irx1b* expression is also lost in vivo (Fig. 7S, W), consistent with our finding that the residual r1 cells in the SU5402 UMAP are molecularly distinct from control r1 cells and display similarity to MHB clusters (Fig. 5E). Expression of both *egr2b* and *mafba* is reduced to a single stripe each (Fig. 6B, F, J, N, Z, DD; Fig. 7B, C, F, G, J, N, R, V). Since simultaneous detection of *egr2b* and *mafba* expression does not reveal an overlap in expression in SU5402-treated embryos (Fig. 7E-H), we conclude that the remaining *egr2b* stripe corresponds to r3 and the remaining *mafba* stripe corresponds to r5 or r6, in agreement with r3 and r6 being partially retained in the SU5402 UMAP (Fig. 5B, Table 1). The fact that *cyp26c1* expression (that marks the forebrain, r2, r5, and r6 at this stage) is completely lost from the hindbrain in SU5402-treated embryos (Fig. 6C, G), further reinforces that the remaining hindbrain clusters in SU5402-treated embryos do not display normal rhombomere identity.

**Figure 7.**
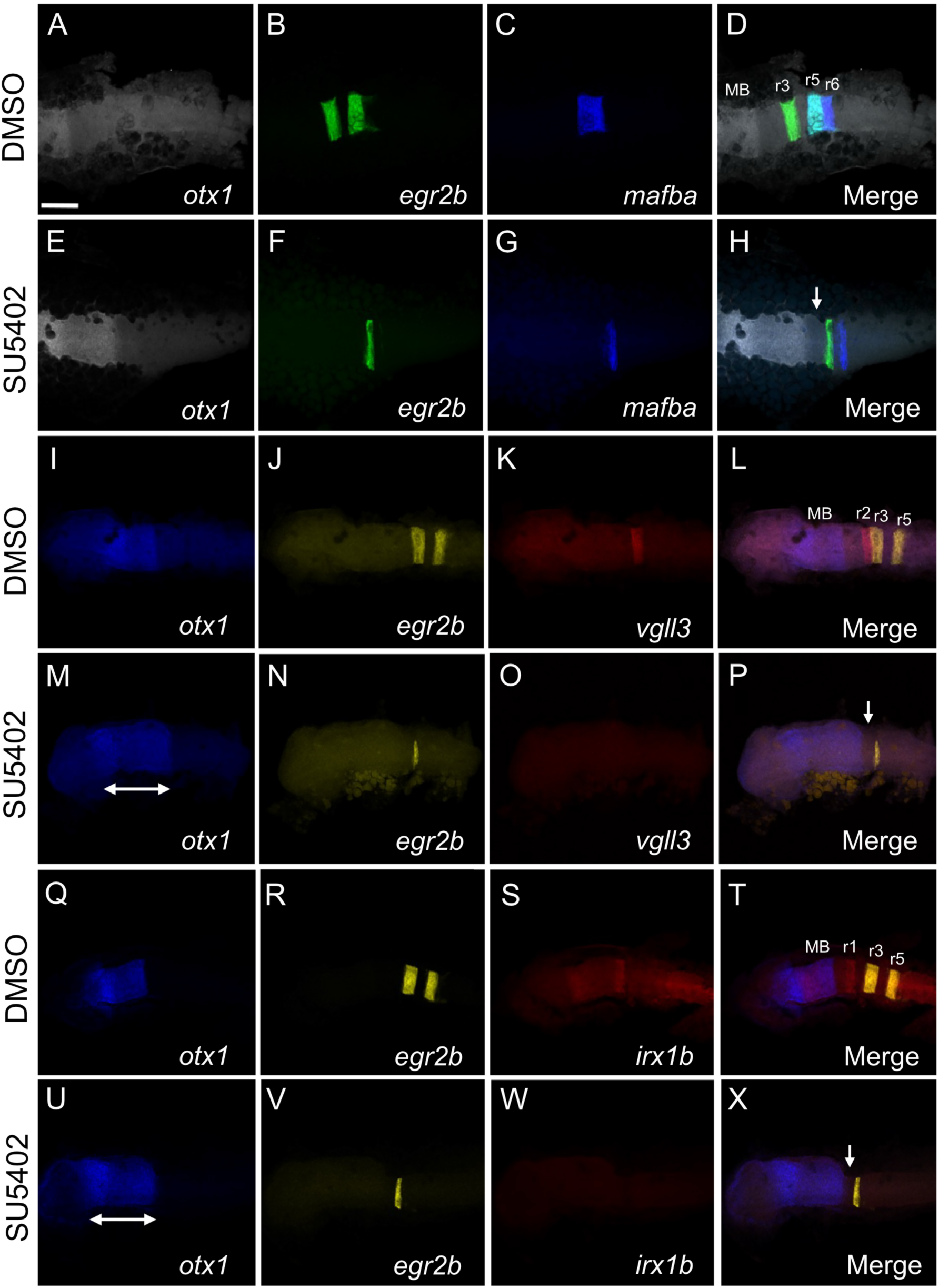
A residual domain is observed in the isthmic region following disruption to Fgf signaling. HCR was used to detect expression of the gene indicated at the bottom of each panel (D, H, L, P, T and X represent merged images of the three panels to their left). Panels show flat-mounted 13hpf DMSO-treated (control) or SU5402-treated, as indicated at left, zebrafish embryos in dorsal view with anterior to the left. Scale bar = 100 um. White arrows in H, P and X indicate gap between *otx1* and *egr2b* expression.

Our analyses indicate that disrupting Fgf-signaling leads to a profound reduction of the isthmic region (including loss of the IsMB, the IsHB, and the IsO), as well as of the hindbrain, although the HCR analyses revealed some residual *egr2b*-positive, and some *mafba*-positive, cells that may correspond to r3 and r5/r6 respectively. These findings are broadly consistent with our scMultiome analysis, which found that the number of cells assigned to the isthmic cell clusters and to the hindbrain clusters were dramatically reduced, but that a few cells remained assigned to these clusters in SU5402-treated embryos – e.g., 5- to 10-fold fewer cells were assigned to each of r1, r3 and r6 in SU5402-treated than in control embryos (Table 1). We note that the expanded *otx1* expression domain in SU5402-treated embryos does not extend to the *egr2b* expression domain but leaves a gap (arrow in Fig. 6FF; 7H, P, X). During normal development, this region between the *otx1* and *egr2b* expression domains corresponds to the isthmic region, r1 and r2. In SU5402-treated embryos, this region lacks expression of all isthmic and hindbrain marker genes we tested by HCR, making its identity unclear. When considered in the context of our scMultiome analysis, this domain is a strong candidate to correspond to the MHB-like and r1 populations, as this would complete the alignment between the cell populations that we identified computationally and those that we observe as anatomically distinct regions in the embryo.

## DISCUSSION

Our study provides a molecular and cellular framework for understanding the organization of the vertebrate isthmus by integrating scMultiomic profiling with spatial validation by HCR in vivo. We identify two major isthmic populations: 1) isthmic midbrain (IsMB), and 2) isthmic hindbrain (IsHB) and further demonstrate that the IsHB itself is subdivided into two distinct subdomains, including an *fgf8*-expressing domain (r0a; likely corresponding to the isthmic organizer – IsO) and a previously uncharacterized *fgf8*-negative domain (r0p). Overall, these findings refine the classical view of the isthmic region and suggest that it is not a singular homogeneous signaling center, but rather a structured and heterogeneous region composed of multiple molecularly distinct cell types.

### Revisiting the organization of the isthmic region

The traditional model of the isthmic region (Fig. 8A) centers on the morphological constriction that it is named after, and that forms at the boundary of the embryonic *Otx* and *Gbx* expression domains (Gibbs et al., 2017; Watson et al., 2018; Puelles 2024). The isthmic organizer is associated with the isthmic region and, after it was found that *fgf8* accounts for organizer activity, the *fgf8* expression domain has generally been taken to define the organizer. While this model has been foundational, it does not account for more recent work suggesting heterogeneity within the isthmic region. Several genes acting at the isthmus (e.g., *Eng* and *Pax* family genes) have expression domains that extend into the adjacent midbrain and hindbrain, suggesting that portions of midbrain and hindbrain located closer to the isthmus may have unique characteristics. Accordingly, prior reports have suggested that r1, which represents the anterior hindbrain, can be further subdivided into r1 and r0 (Vaage 1969; Moens and Prince, 2002; Zhi-Rong et al., 2010). Also, since *fgf8* is expressed in the anterior hindbrain, it has been suggested that the IsO may represent a distinct domain of r1 (Puelles and Rubenstein, 2015; Puelles, 2024). While compelling, these proposed subdivisions of the isthmic region are based on the expression of only a few genes, and it is not clear if they refer to the same domains – e.g., is the proposed r0 identical to the proposed IsO? Using scMultiome, which provides comprehensive transcriptional and chromatin accessibility data at single-cell resolution, we have expanded on this prior work by molecularly defining cell populations present at the isthmus. We find that cells within the isthmus segregate into two transcriptionally distinct populations aligned along the anterior-posterior axis (Fig. 8B). The IsMB exhibits a hybrid identity, co-expressing isthmic markers (e.g., *en2b*, *pax2a*) alongside midbrain-associated genes such as *otx1* and *wnt1*, making it distinct from other midbrain populations. In an analogous fashion, the IsHB population is defined by co-expression of isthmic markers with hindbrain-associated ones, including *gbx2* and *fgf8*, making this domain molecularly distinct from the rest of the anterior hindbrain. Based on its gene expression profile, we propose that the IsHB corresponds to the previously postulated r0 domain.

**Figure 8.**
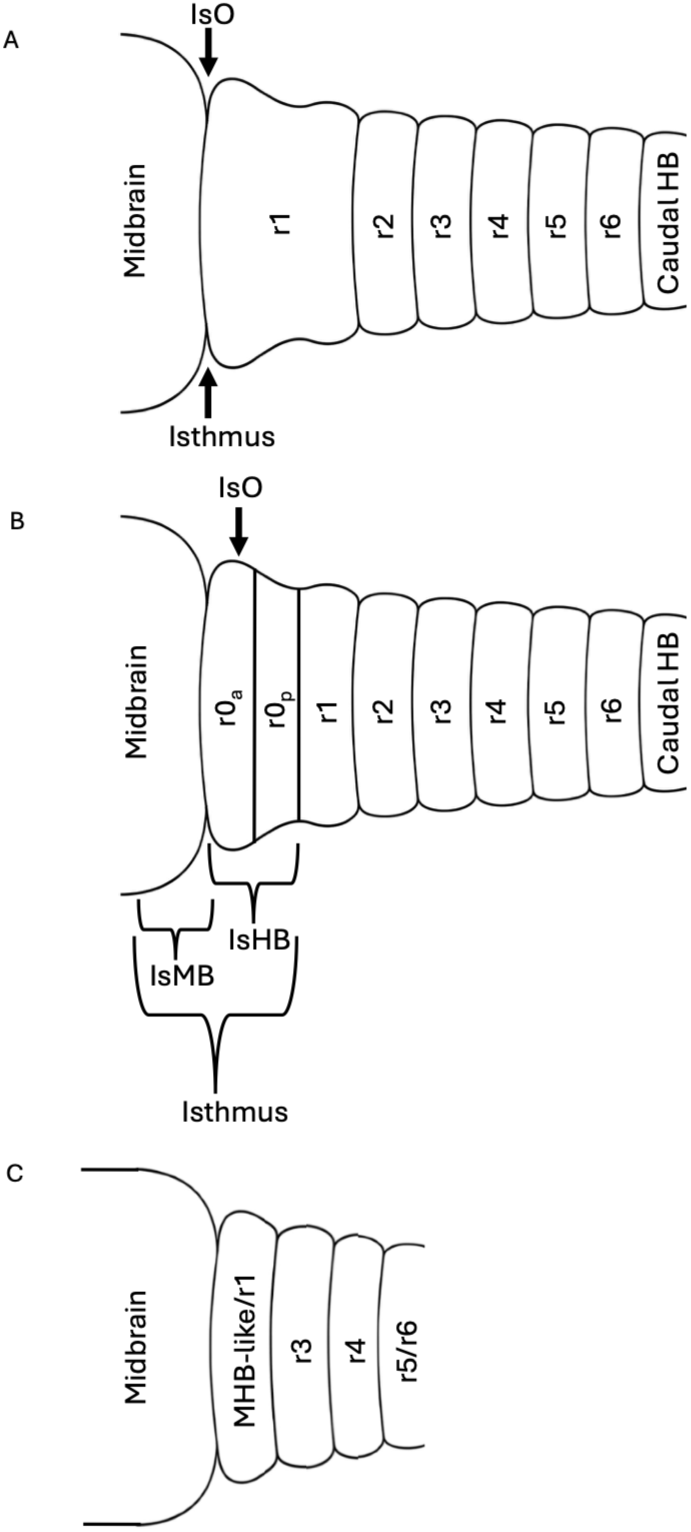
Schematic diagrams of the isthmus region. (A) Traditional model of the vertebrate isthmus. (B) Proposed model of the vertebrate isthmus. (C) Proposed model of the SU5402-treated vertebrate isthmus.

Within the IsHB/r0, we identified two subpopulations distinguished primarily by *fgf8* expression (Fig. 8B). Since *fgf8* marks the IsO, this means that the IsO is not identical to r0. Instead IsHB/r0 consist of two subdomains (r0a and r0p) where the anterior r0a domain expresses *fgf8*, and thus corresponds to the IsO, and the posterior r0p domain that lacks *fgf8* expression represents a previously undescribed domain. Spatial HCR validation confirms that these subpopulations are physically segregated during brain development in vivo, with r0p positioned between IsO/r0a and r1. Our work provides molecular support for the long-debated concept of a distinct anterior hindbrain segment (IsHB/r0), while also demonstrating that IsHB/r0 itself is heterogeneous rather than a single entity. The identification of IsO/r0a and r0p as molecularly distinct subdomains suggests that the region anterior to r1 is more complex than previously appreciated. Rather than only containing the organizer, this region may be comprised of multiple functionally distinct compartments, with potential specialized roles in patterning and signaling. Whether r0p contributes directly to organizer activity, functions as a transitional territory between the IsO and r1 or represents a distinct anterior hindbrain subdomain remains to be determined.

### Early establishment of isthmic heterogeneity

We find that the IsMB and IsHB populations are already molecularly defined at 13hpf – prior to the isthmus being identifiable morphologically – indicating that isthmic heterogeneity emerges early during brain development rather than being a later refinement. While the IsHB subdomains are detectable at this stage, the *fgf8* signal is less robust, perhaps reflecting progressive maturation of the isthmic region. These results suggest that subdivision of the isthmus is an intrinsic feature of its developmental program and may be established concurrently with, or shortly after, the initial positioning of the Otx/Gbx boundary. The presence of distinct chromatin accessibility signatures and enriched transcription factor motifs in each population further supports the idea that these subdomains are governed by distinct gene regulatory networks. For example, enrichment of OTX-related motifs in IsMB and hindbrain-associated motifs in IsHB indicates that regional identity is tightly coupled to underlying regulatory architecture.

### Fgf signaling is required for isthmic subdivision and identity

Our experiments using the Fgf inhibitor SU5402 demonstrated that Fgf signaling is essential for the establishment of the isthmic subdomains. Upon inhibiting Fgf signaling, we observe a disruption of isthmic heterogeneity, with the loss of both IsMB and IsHB populations and the persistence of only a single isthmus-related cluster (MHB.2 or MHB-like; Fig. 8C). This is accompanied by the loss of key isthmic markers (*fgf8*, *wnt1*, *pax2a*, *en2b*) and a dramatic reduction in hindbrain rhombomeres, consistent with previous studies (Reifers et al., 1998; Chi et al., 2003) demonstrating the central role of Fgf signaling in hindbrain patterning.

Interestingly, the remaining SU5402-associated cluster retains transcriptional similarity to the control MHB.2 population, suggesting that it may represent a residual or partially specified isthmic-like state. We propose that this cluster may reflect an amalgamated identity, arising from cells that would normally segregate into IsMB, r0a, r0p, and possibly r1, but fail to do so in the absence of Fgf signaling (Fig. 8C). This interpretation is supported by the observed expansion of anterior (midbrain) identity and the presence of a gap between midbrain and remaining r3 structure in SU5402-treated embryos. These findings reinforce a model in which Fgf signaling is not only required for organizer function but also for cellular diversification within the isthmus, and surrounding architecture (e.g., r1), potentially by stabilizing distinct transcriptional states or reinforcing positional identity along the anterior-posterior axis.

### Limitations and future directions

While our study provides strong evidence for isthmic heterogeneity, several questions remain. Firstly, the functional role of the r0p population is unknown. It will be important to determine whether these cells contribute to organizer signaling, act as a transitional buffer zone, or represent a lineage-restricted precursor population. Secondly, fate mapping and perturbation experiments will be required to establish the developmental trajectories and fate potential of IsMB, r0a, and r0p cells, especially at earlier developmental timepoints. Thirdly, we have not tested the role for other signaling pathways, such as Wnt and retinoic acid signaling, and these may also contribute to the establishment of isthmic domains. The SU5402 inhibitor we use must diffuse into the embryo and we may not have completely blocked all Fgf signaling. Finally, extending this analysis to later developmental stages and to other organisms will be important for determining the extent to which these findings are conserved and how early isthmic heterogeneity affects brain development.

## Conclusion

In summary, our work reveals that the vertebrate isthmus is composed of multiple transcriptionally and spatially distinct subpopulations that arise early during development and depend on Fgf signaling for their formation. By redefining the isthmus as a heterogeneous domain, our study builds on prior work to provide a conceptual framework for understanding how organizer regions function to pattern the developing brain.

## EXPERIMENTAL PROCEDURES

### Zebrafish husbandry and embryo collection

All zebrafish procedures were approved by the Institutional Animal Care and Use Committee (IACUC) at the University of Colorado School of Medicine under protocol #870. Wild-type TU (ZL57) zebrafish were obtained from the Zebrafish International Resource Center and reared in our facility. For embryo collection, one adult male and one adult female zebrafish were separated overnight in breeding tanks and paired the following morning. Eggs were collected in 10 cm Petri dishes, maintained in egg water containing 60 µg/mL Instant Ocean and 0.0002% methylene blue, and incubated at 29°C until the desired developmental stage. Dead and unfertilized eggs were manually removed and excluded from analyses.

### FGF signaling inhibition with SU5402

SU5402 (Abcam, ab141368) was used to inhibit Fgf signaling. SU5402 was dissolved in DMSO and diluted to a final concentration of 10 µM. Embryos were randomly assigned to DMSO control or SU5402 treatment groups and treated beginning at 7hpf until collection at 13hpf. Embryos were dissected using forceps (Fine Science Tools Dumont #5 Forceps, Cat #NC9889584) to exclude yolk and posterior tissues. DMSO-treated embryos were used as controls. Dead embryos were manually removed and excluded from our analyses.

### Single-cell multiome (scMultiome) sample preparation and analysis

Single-cell multiome RNA-seq and ATAC-seq data from 13 and 16 hours-post-fertilization (hpf) zebrafish embryos were generated and processed as described previously in Kim et al., 2023. Twenty-five SU5402-treated embryos were collected at 13hpf and dissected using forceps (Fine Science Tools Dumont #5 Forceps, Cat #NC9889584) to include the entirety of the midbrain and hindbrain. Dead embryos were manually removed and excluded from our analyses. Tissue was collected in 1X PBS, mechanically dissociated by repeated pipetting through a P1000 tip, and centrifuged at 400 × g for 5 min at 4°C. The resulting pellet was resuspended in 500 μl of protease solution containing 10 mg/ml BI protease (Sigma, P5380), 125 U/ml DNase, and 2.5 mM EDTA in PBS, and incubated on ice for 15 min. Samples were then centrifuged again at 400 × g for 5 min at 4°C, resuspended in 1 ml HBSS supplemented with 2% FBS, filtered through a 20 μm cell strainer (pluriSelect, KL-071912), and centrifuged once more under the same conditions. The pellet was resuspended in 500 μl HBSS + 2% FBS and passed through a second 20 μm cell strainer. After a final centrifugation at 400 × g for 5 min at 4°C, cells were resuspended in 200 μl PBS.

For nuclei isolation, the 10X Genomics recommended protocol was followed with minor modifications. Dissociated cells were centrifuged at 900 × g for 5 min at 4°C, resuspended in 100 μl of 0.1X lysis buffer consisting of 1 mM Tris-HCl pH 7.4, 1 mM NaCl, 0.3 mM MgCl₂, 0.1% BSA, 0.01% Tween-20, 0.01% NP-40, 0.001% Digitonin (Invitrogen, BN2006), 0.1 mM DTT, and 0.1 U/μl RNase inhibitor in nuclease-free water, and incubated on ice for 5 min. Samples were subsequently washed three times by centrifugation at 900 × g for 10 min at 4°C followed by resuspension in 1 ml chilled wash buffer containing 10 mM Tris-HCl pH 7.4, 10 mM NaCl, 3 mM MgCl₂, 1% BSA, 0.1% Tween-20, 1 mM DTT, and 1 U/μl RNase inhibitor in nuclease-free water. Nuclei were then counted and resuspended in 1X nuclei buffer provided with the 10X Genomics Single-Cell Multiome ATAC Kit A, supplemented with 1 mM DTT and 1 U/μl RNase inhibitor in nuclease-free water, to a final concentration of approximately 2,000–3,000 nuclei/μl for sequencing on the 10X scMultiome platform.

Fastq sequencing files from 10X Genomics single-cell multiome RNA-seq and ATAC-seq libraries were processed using Cell Ranger ARC v1.0.1 with the zebrafish GRCz11 reference library to generate UMI-based gene expression counts and ATAC peak fragment counts. Data were analyzed in R v.4.5.3 using standard workflows from the Signac v1.16.0 package. Briefly, Seurat objects were generated from the ‘matrix.h5’ and ‘fragments.tsv.gz’ files, annotated using GRCz11.v99.EnsDb, and ATAC peaks were corrected by peak calling with MACS2 v2.2.7.1. Cells were then filtered to remove low-quality cells with low total RNA expression, low total ATAC counts, high mitochondrial gene expression, nucleosome signal >2, or TSS enrichment <1. Gene expression data were normalized using SCTransform, followed by dimensionality reduction with PCA. DNA accessibility data were processed using latent semantic indexing. A weighted nearest neighbor graph was generated using Seurat and visualized by UMAP, and clusters were annotated based on the expression of known marker genes. Per-cell motif activities were scored using chromVAR. Significantly differentially expressed genes and differentially active motifs were identified for each cell cluster.

To integrate Seurat objects, the full intersecting set of ATAC peaks containing peaks from any of the three datasets was identified, and a new chromatin accessibility assay was generated for each object using the full intersecting peak file. RNA-seq data were integrated using the Seurat v5 integration workflow, while the new chromatin accessibility data were integrated using Harmony. The integrated RNA and ATAC datasets were then analyzed with the Seurat weighted nearest neighbor method to compute UMAP embeddings and identify clusters.

Cell types and UMAP embeddings for the SU5402-treated 13hpf Seurat object were predicted using the untreated control 13hpf Seurat object as a reference, following the Seurat v5 multimodal reference mapping tutorial. For the SU5402-treated dataset, cells were retained without additional filtering to preserve the maximum number of cells from the sample. Cell counts for clusters expressing a given gene were calculated based on cells with normalized expression values greater than 0 for that gene.

Clusters were annotated based on expression of known regional markers for forebrain, midbrain, isthmus, and hindbrain identities. Isthmic populations were identified based on expression of canonical markers including *en2b*, *pax2a*, *pax5*, *wnt1*, *fgf8a*, *fgf8b*, and *gbx2*. Heatmaps comparing isthmic clusters with midbrain and r1 populations, as well as heatmaps comparing SU5402-treated isthmic clusters with control isthmic clusters, were generated using dittoHeatmap. Differentially expressed gene (DEG) analysis was performed using FindAllMarkers, and genes were considered significant using an adjusted p-value adjusted threshold of ‘0.05’ and log2 fold-change cutoff of ‘0.25’. Motif enrichment analysis was performed using Signac v.1.16.0/chromVAR/JASPAR2020 database, and enriched transcription factor motifs were visualized for each isthmic population. Dot plots were generated with DotPlot. Volcano plots were generated with EnhancedVolcano v1.28.2, with genes considered significant at p < 0.05. Gene Ontology (GO) Biological Process enrichment analysis was performed using clusterProfiler v4.18.4, with zebrafish gene annotations from org.Dr.eg.db v3.22.0; significantly enriched GO terms were identified using adjusted p-values threshold of ’0.05’. Schematics were generated using Adobe Photoshop 2024 (version 24.7.0). Similarity between SU5402-associated MHB clusters and control clusters was evaluated using two complementary methods: module score analysis based on the SU5402 MHB.2 gene signature and cluster similarity analysis with ClusterFoldSimilarity v1.0.0. Module scores were calculated using AddModuleScore, and similarity scores were displayed as heatmaps. scMultiome data has been deposited in GEO under record number GSE337538 and the code used to generate the data for each figure is available at GitHub at https://github.com/rebeccaorourke-cu/Nunez_MBHB_manuscript/.

### Hybridization chain reaction RNA FISH analysis

In vivo gene expression was detected by hybridization chain reaction (HCR). HCR RNA probe bundles were purchased from Molecular Instruments (https://www.molecularinstruments.com) for the following probes: *en2b*, *nr2f2*, *otx1*, *pax2a*, *wnt1*, *gbx2*, *her5*, *ism1*, *en2a*, *fgf8a*, *egr2b*, *cyp26c1*, *mafba*, *vgll3*, *irx1b*. Embryos were fixed in 4% paraformaldehyde, dehydrated in 100% methanol, and stored at −20°C until further use. Prior to hybridization, embryos were rehydrated through a graded methanol/PBST series, with each wash performed for 5 min: 75% methanol/25% PBST, 50% methanol/50% PBST, 25% methanol/75% PBST, and 100% PBST. PBST consisted of 1X PBS with 0.1% Tween-20. Embryos were pre-hybridized in probe hybridization buffer provided by the manufacturer for 30 min at 37°C, followed by overnight incubation at 37°C in hybridization buffer containing the desired HCR probe sets, with 2 pmol of each probe set. The following day, the probe solution was removed, and embryos were washed four times for 15 min at 37°C using manufacturer-provided probe wash buffer, followed by two 5 min washes in 5X SSCT, consisting of 5X SSC with 0.1% Tween-20. Embryos were then pre-treated in amplification buffer for 30 min at room temperature. Separately, h1 and h2 hairpins were prepared by snap cooling a 3 μM stock: hairpins were heated to 95°C for 90 s and then cooled to room temperature in a dark drawer for 30 min. The h1 and h2 hairpins were added to amplification buffer and incubated with embryos overnight in the dark at room temperature. The next day, the hairpin solution was removed, embryos were washed with 5X SSCT, and samples were stored at 4°C until imaging.

HCR analysis to confirm the expression pattern of published genes was carried out on a minimum of 5 embryos and 2 biological replicates. The number of embryos assayed by HCR for overlap of gene expression in Figure 2A-V were as follows: *en2b*/*nr2f2* – 5 embryos, two biological replicates; *en2b*/*otx1* – 5 embryos, two biological replicates; *pax2a*/*wnt1* – 5 embryos, two biological replicates; *en2b*/*gbx2* – 5 embryos, two biological replicates; en2b/fgf8a – 5 embryos, two biological replicates; *fgf8a*/*nr2f2* – 5 embryos, two biological replicates; *en2a*/*her5*/*ism1* – 5 embryos, two biological replicates.

The number of embryos assayed by HCR for overlap of gene expression in Figure 4A-R were as follows: *en2b*/*nr2f2* – 5 embryos, two biological replicates; *en2b*/*otx1* – 5 embryos, two biological replicates; *pax2a*/*wnt1* – 5 embryos, two biological replicates; *en2b*/*gbx2* – 5 embryos, two biological replicates; *en2b*/*fgf8a* – 5 embryos, two biological replicates; *fgf8a*/*nr2f2* – 5 embryos, two biological replicates.

The number of SU5402-treated embryos assayed by HCR for overlap of gene expression in Figure 6A-FF were as follows: *pax2a*/*egr2b*/*cyp26c1* – 10 embryos, three biological replicates; *en2b*/*egr2b*/*fgf8a* – 10 embryos, three biological replicates; *otx1*/*gbx2*/*fgf8a* – 10 embryos, three biological replicates; *otx1*/*egr2b*/*wnt1* – 10 embryos, three biological replicates.

The number of SU5402-treated embryos assayed by HCR for overlap of gene expression in Figure 7A-X were as follows: *otx1*/*egr2b*/*mafba* – 10 embryos, three biological replicates; *otx1*/*egr2b*/*vgll3* – 10 embryos, three biological replicates; *otx1*/*egr2b*/*irx1b* – 10 embryos, three biological replicates.

An equivalent number of control embryos treated with DMSO (vehicle control) was used. All drug-treated embryos displayed minor variations in the phenotypes shown in Figures 6A-FF and 7A-X, and all vehicle-treated embryos displayed wildtype expression. Sample size was determined based on pilot experiments which demonstrated that all treated embryos showed the reported phenotype. Biological replicate is taken to mean that each replicate was performed on a set of embryos that were treated separately throughout the experiment. The scoring of drug-treated embryos was not blinded.

### Embryo preparation for microscopy and confocal imaging

After being fixed and stained for HCR, embryos were microdissected to remove the yolk and posterior body (somites and tailbud) using forceps (Fine Science Tools Dumont #5 Forceps, Cat#NC9889584). Dissected embryos were flat-mounted onto a microscope slide (Fisherbrand™ Frosted Microscope Slides, Cat#12-550-400), embedded in 20ul of Fluoromount-G (SouthernBiotech, Cat#0100-01) and covered with a glass coverslip (BoliOptics, Cat#SL39201012) prior to imaging. Laser scanning confocal microscopy was conducted on a Zeiss LSM880. Images were collected with a x10/0.8 air-objective lens. All channels were captured sequentially with minimal speed in bidirectional mode, with the range of detection attuned to avoid overlap between channels. Image processing was conducted using ZEN (Zeiss, version 3.12) and followed the same settings throughout. Acquired Z-stacks were saved, cropped and rotated using Adobe Photoshop 2024 (version 24.7.0).

## Author contributions

Conceptualization: S.A.N., Y.I.K., R.O., C.G.S.; Investigation: S.A.N.; Formal analysis: S.A.N., R.O.; Resources: C.G.S.; Writing - original draft: S.A.N.; Writing - review & editing: R.O., C.G.S.; Supervision: C.G.S.; Project administration: C.G.S; Funding acquisition: S.A.N., C.G.S.

## Data availability

scMultiome data has been deposited in GEO under record number GSE337538

## Code availability

The code used to generate the data for each figure is available at GitHub at https://github.com/rebeccaorourke-cu/Nunez_MBHB_manuscript/.

